# Hyperglycemic state and fluid shear stress affect metastatic breast cancer cell migration via focal adhesion kinase

**DOI:** 10.1101/2025.05.22.655615

**Authors:** Brandon D. Riehl, Eunju Kim, Taylor Boudreaux, Osaira Ovando, Stephanie Vielmas- Duarte, Suyong Choi, Hamid Band, Lanju Mei, Diganta Dutta, Surabhi Chandra, Jung Yul Lim

## Abstract

We tested how the diabetes-related hyperglycemic condition affects the migration of highly metastatic triple-negative breast cancer (TNBC) cells, MDA-MB-231, under a physiological fluid shear environment. MDA-MB-231 cells displayed a significantly enhanced migratory behavior under a high glucose condition (25 mM) specifically when exposed to flow at 15 dyne/cm^2^ shear stress. In contrast, the effect of fluid shear was marginal under low glucose (5 mM). Normal epithelial MCF-10A cells, on the other hand, showed increased migration by fluid shear under both low and high glucose conditions. The fluid shear-triggered MDA-MB-231 cell migration under high glucose was significantly abrogated by a focal adhesion kinase (FAK) inhibitor, supporting the mediatory role of FAK in MDA-MB-231 TNBC cell sensing of the high glucose-fluid shear environment during migration. The role of FAK was further demonstrated by the effects of FAK inhibitor on MDA-MB-231 cell migration in scratch wound healing and Boyden chamber migration assays. Our studies provide evidence that high glucose and fluid shear could jointly trigger MDA-MB-231 TNBC cell migration that requires FAK activity. These may provide improved mechanistic insights into how concurrent diabetes may impact the pro-metastatic behavior of breast cancer and suggest the impact of exploring FAK as a relevant therapeutic target.

---

Metastatic disease accounts for nearly 90% of cancer deaths (United States Cancer Statistics data [1]). It is estimated that metastatic breast cancer has a 5-year survival rate of about 30% in the US. Breast cancer patients with coexistent diabetes exhibit poor prognosis, chemoresistance, increased metastasis, and a 40% increased risk of mortality compared to women without diabetes [2]. Triple-negative breast cancer (TNBC), in which the tumor cells lack the receptors for estrogen receptor, progesterone receptor, and HER2 protein [3], is an aggressive subgroup of breast cancer that is highly invasive. Current treatment options for TNBC are mostly limited to conventional chemotherapy and radiation as targeted therapies are lacking, and the rates of recurrence and progression remain high. As a result, the median survival rate of metastatic TNBC patients is approximately 10.2 months, while it is about 5 years for non-TNBC breast cancer patients [4].

Metastasis is a complex process involving the detachment of cancer cells from the basement membrane, leading to cell migration and invasion, extravasation, and growth of secondary tumors. A key component of cancer cell migration involves the remodeling of the cancer cell cytoskeleton, including actin, microtubule, and intermediate filaments [5]. Directed cell migration occurs in response to gradients of growth factors or chemokines as well as to the mechanophysical cues from the extracellular matrix (ECM) [6]. Cell migration occurs through protrusion, adhesion, contraction, and retraction, and actin cytoskeletal architecture and focal adhesion play a key role in executing cell movement [7]. Cell membrane protrusions driven by actin polymerization are extended towards biochemical and mechanophysical extracellular cues. New focal adhesions stabilize these protrusions by linking the actin cytoskeleton to ECM proteins. Actomyosin contraction promotes the disassembly of these adhesions at the rear end of the cell. Retraction then brings the trailing cell body forward in the direction of cell movement to complete the migration process.

Cytoskeletal regulatory proteins such as RhoA kinase (ROCK) and focal adhesion proteins such as focal adhesion kinase (FAK) assist in the migration of cells. ROCK regulates the activation of multiple actin filament formation axis proteins (LIMK, cofilin, etc.), and coordinates with non-muscle myosin II to generate the contractile force to drive motility [8]. FAK regulates the response of cells to the environmental stimuli from ECM by sensing changes in extracellular mechanical forces and controlling actin remodeling during focal adhesion formation [9]. Focal adhesions link ECM proteins to actin cytoskeleton via linker proteins including talin, vinculin, paxillin, and FAK, with the balance between adhesion and contraction, thereby modulating cell migration [10]. The loss and formation of focal adhesions act in the separation from the basement membrane and new focal contact establishment at the new matrix site, causing cell migration.

FAK has been implicated in the progression of breast cancer, and several recent attempts explored FAK inhibitors as antimetastatic agents. For example, immunotherapy using cytokine-induced killer (CIK) in combination with a FAK inhibitor was found to induce higher TNBC cell death than CIK treatment alone [11]. An antihistamine drug, Ebastine, was found to reduce tumors via blocking the tyrosine kinase domain of FAK and thereby downregulating angiogenesis [12]. FAK inhibition combined with retinoids (derivatives of retinoic acid from Vitamin A) could also be effective at inhibiting TNBC tumorigenesis and metastasis in vivo [13]. Connexin 31-mediated tumor cell homotypic adhesion and tumor-astrocyte heterotypic adhesion were shown to function through FAK activation to promote TNBC brain metastasis, and systemic treatment with FAK inhibitors could reduce brain metastasis progression [14]. Thus, substantial literature supports a role of FAK in TNBC metastasis, proposing FAK as a therapeutic target.

An association between type 2 diabetes mellitus (T2DM) and breast cancer mortality has been proposed [15,16], but underlying mechanisms are not fully known. Recently, it was shown that RAGE (receptor for advanced glycation end products) is elevated in TNBC and promotes metastasis through the activation of FAK-Hippo/YAP pathway [17]. Considering that the AGE-RAGE axis has been extensively targeted for diabetes [18–20], combined data suggest a potential that the regulatory cascade of RAGE such as FAK could play a role in the cancer-diabetes interaction. In addition to its role in cancer, FAK is recently proposed to be implicated in promoting diabetic cardiomyopathy as increased FAK phosphorylation was found in the hearts of streptozotocin-induced type 1 diabetic mice [21] and contributed to arterial calcification [22].

While studies support the potential role of FAK in linking the association of T2DM and breast cancer prognosis and metastasis, there is a gap in knowledge regarding how diabetes promotes the basic pro-metastatic cellular processes. We have previously demonstrated that highly metastatic TNBC cells (MDA-MB-231) show stimulated cell migration under physiologically relevant fluid shear stress environment in comparison with less metastatic MDA-MB-468 breast cancer cells and non-tumorigenic MCF-10A epithelial cell control [23]. However, the underlying molecular mechanosensing mechanisms remain unclear, and how diabetes-relevant conditions might impact these behaviors is not known. In this study, we tested the hypothesis that FAK is involved in promoting the migratory behavior of TNBC cells triggered by diabetes-related hyperglycemic state in conjunction with physiologically relevant fluid shear stress condition. Considering FAK inhibitors as a potential antimetastatic agent, we pursue the question of whether metastasis capability is restored by the shear stress and hyperglycemic conditions, or if FAK inhibition is sufficient on its own to abrogate the effects of flow and glucose. Our approach measures the functional effect of fluid shear, hyperglycemia, and FAK inhibition on detailed and nuanced metastasis parameters and invasion assays. We show, for the first time to our best knowledge, that a high glucose condition mimicking T2DM-associated hyperglycemia collaborates with fluid shear stress to promote MDA-MB-231 TNBC cell migration through the activation of FAK.

## Results

### Non-metastatic normal MCF-10A epithelial cell migration shows dependency on glucose level and fluid shear

We first conducted tests on the effects of glucose concentration and fluid shear on cell migration using non-tumorigenic MCF-10A epithelial cells. Co-centered individual cell migration tracks are plotted in distinct colors (Fig. 1a) and as vector plots showing starting and ending points (Fig. 1b). These plots provide observations of the overall migration trend of each group including the migration paths and the tendency for directed migration. MCF-10A cells responded to both glucose concentration and fluid shear. Cells under low glucose (marked as L, 5 mM) condition with no flow (static) had the least cell migration displacement (Fig. 1c). Exposure to high glucose (marked as H, 25 mM) and fluid shear stress at 15 dyne/cm^2^ (FF15) significantly increased cell displacement. The cell migration distance under high glucose static condition was comparable to that under low glucose with shear flow. The confinement ratio, a measure of the directness of the cell path, was increased by high glucose both under static and flow conditions (Fig. 1d). The confinement ratio determines whether cells wander randomly and stay close to the point of origin, or if the cells move towards a directed goal with little deviation from the path. The arrest coefficient, a measure of pausing during migration, was not affected by change in glucose concentration but was significantly decreased under fluid shear (Fig. 1e). The arrest coefficient can distinguish, for example, between a cell that steadily migrates to a destination, and a cell that covers the same distance faster but later lays fallow. Collectively, the results using normal MCF-10A epithelial cells showed an increased cell migration under high glucose and fluid shear. This could be achieved by MCF-10A cells traveling in a more direct path under high glucose and pausing less under fluid shear stress. The trends of overall migration of cells (i.e. the ensemble-average of all-participating cells) are shown in the root-mean-square (RMS) displacement plots (Fig. 1f). The high glucose and fluid shear group had the largest RMS displacement, indicating that high glucose and shear effects work together to promote the migration of normal MCF-10A mammary epithelial cells. The motility coefficient, the measure of migration strength quantified from the slope of the RMS plot similar to the measure of diffusion coefficient, displayed similar trends with respect to glucose and shear (motility coefficient values noted in Fig. 1 caption).

**Fig. 1.**
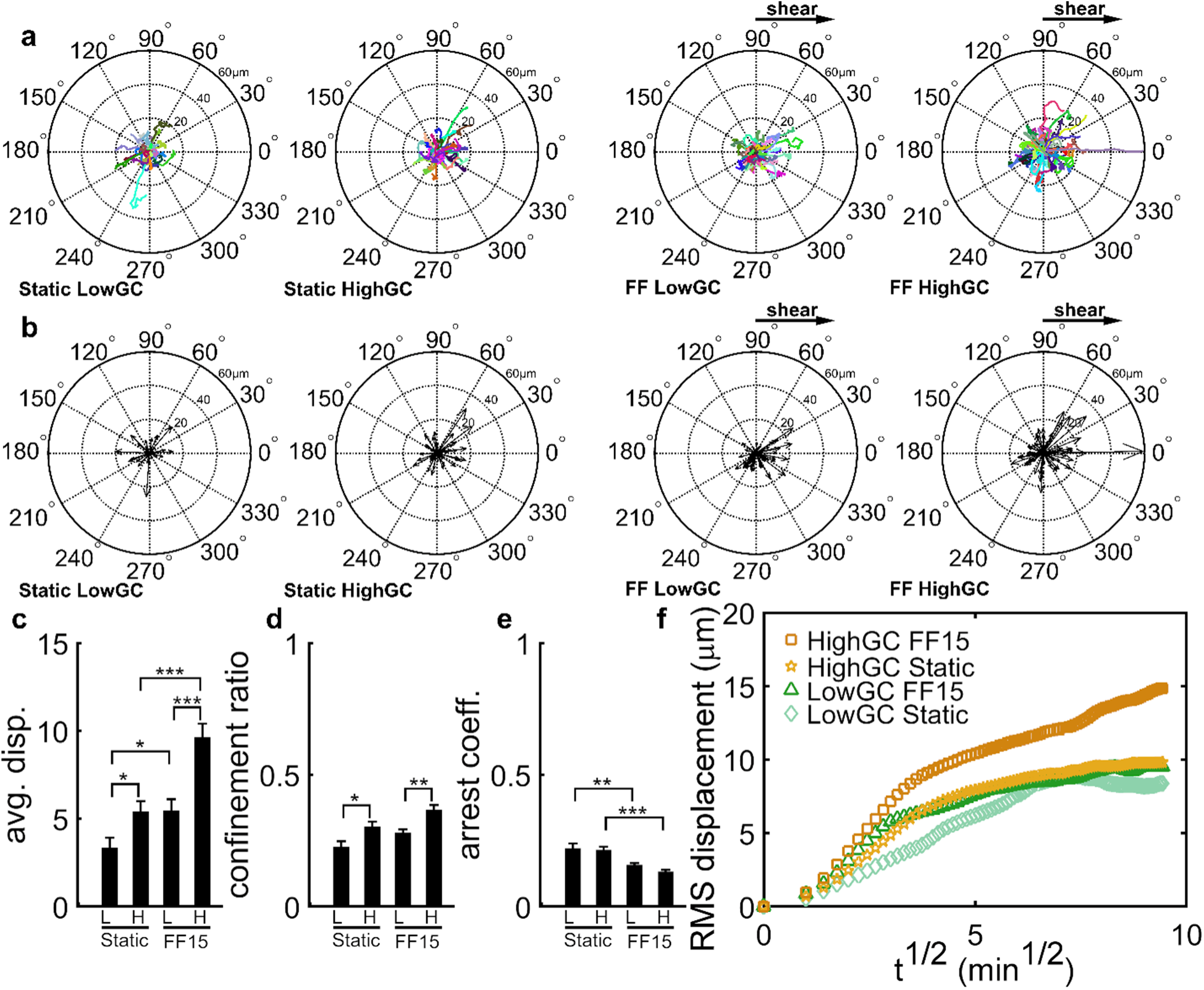
MCF-10A normal epithelial cell migration is induced by fluid shear and media glucose level. The fluid shear direction is from left to right. Co-centered migration tracks (a) and vector plots (b) show individual cell migration tracks. The static (no flow) and fluid flow (FF) conditions with low and high glucose concentration (GC) are shown. The average cell displacement (c) increased with high glucose (labeled: H) and with fluid shear (FF15: 15 dyne/cm^2^). Cells in the high glucose condition tended to migrate in more direct paths as measured by the confinement ratio compared to the low glucose condition (labeled: L) (d). Cells in the fluid sheared condition migrated with fewer pauses for both low and high glucose levels, as measured by the arrest coefficient (e). As an overall measure of ensemble-averaged cell migration, the RMS displacement is plotted (f). Motility coefficient (slope of RMS displacement vs. t^1/2^) was LowGC Static: 0.93, HighGC Static: 0.91, LowGC FF15: 0.81, and HighGC FF15: 1.35. *, **, ***: p < 0.05, 0.01, and 0.001, respectively. Bar graphs are plotted as mean ± SEM.

### Highly metastatic MDA-MB-231 TNBC cells exhibit enhanced migration under high glucose when exposed to fluid shear

Using the glucose concentration and fluid shear setups described above, we obtained time lapse images of MDA-MB-231 TNBC cell migrations (Fig. 2a,b). Unlike MCF-10A cells, MDA-MB-231 cells under static condition had a lower average migration and a lower confinement ratio in the high glucose condition compared to low glucose (Fig. 2c,d). A difference in the arrest coefficient between high and low glucose was not observed under static condition (Fig. 2e). Intriguingly, shear flow stimulated the migration of metastatic MDA-MB-231 cells especially for the case of high glucose situation, i.e., MDA-MB-231 cells migrated further (Fig. 2c), in more direct paths (Fig. 2d), and with fewer pauses (Fig. 2e) under high glucose and fluid shear. Migration along the flow direction was also increased for MDA-MB-231 cells with high glucose and fluid shear (Fig. 2f), which was assessed as the percentage of cells migrating within π/4 of the fluid flow direction (Fig. 2g). Cells in low glucose under fluid shear also showed an increased trend to migrate along the flow direction but to a lesser extent. Overall, the combined effects of high glucose and fluid shear were to promote the migration of MDA-MB-231 TNBC cells.

**Fig. 2.**
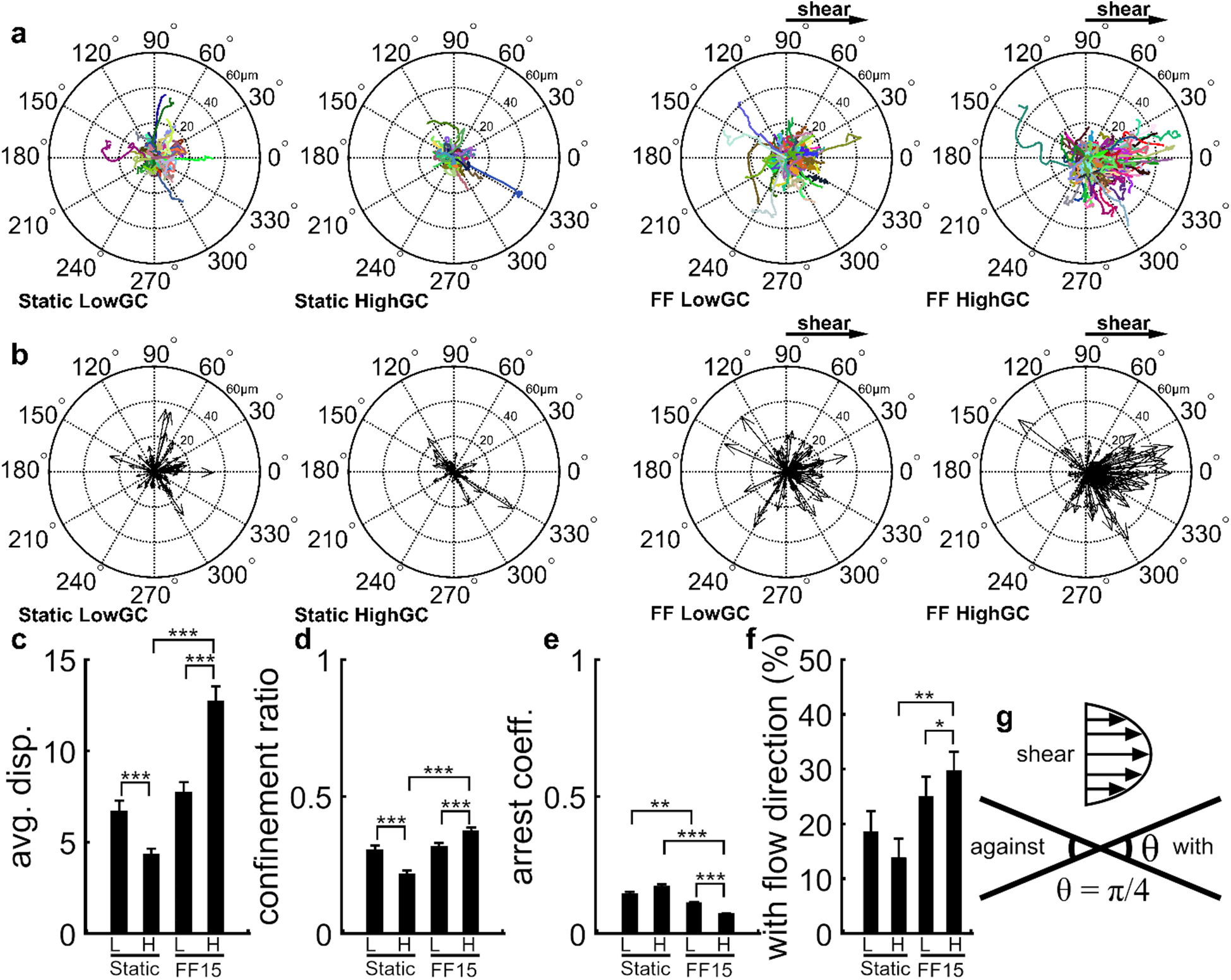
MDA-MB-231 triple negative breast cancer cells have high inducibility to directed migration via combined high glucose and fluid shear. Co-centered cell migration tracks (a) and corresponding vector plots (b). The average MDA-MB-231 cell displacement (c) and confinement ratio (d) were significantly increased by the combined effects of high glucose (labeled: H) and fluid shear (FF15: 15 dyne/cm^2^). The effects of fluid shear were minimal under low glucose (labeled: L). MDA-MB-231 cells paused less during migration when exposed to fluid shear in high glucose (e). More cells migrated along the flow direction under high glucose and fluid shear (f), as assessed for the cells maintaining migration path within π/4 of the flow direction (g). *, **, ***: p < 0.05, 0.01, and 0.001, respectively. Bar graphs are plotted as mean ± SEM.

The increased migration capability of MDA-MB-231 cells in the high glucose and fluid shear combined condition was prominent in the cell migration speed. At the shear onset, cells under shear flow in both low and high glucose conditions had higher migration speeds compared to static controls (Fig. 3a). Cells under high glucose and exposed to fluid shear had the highest migration speeds, almost twice that of the other conditions. The higher migration speed persisted for more than 30 min after the flow onset. Collectively, the sustained higher migration speed, more efficient migration path, and fewer stops resulted in a much higher RMS group displacement for the high glucose plus shear flow condition (Fig. 3b). MDA-MB-231 cells in low glucose and exposed to fluid shear also had increased RMS displacement compared to the static condition, but the difference was less. The motility coefficient (noted in Fig. 3 caption) represented these changes. The combined results indicate that highly metastatic MDA-MB-231 TNBC cells become significantly more mobile when encountering fluid shear stress and diabetes-relevant high glucose condition.

**Fig. 3.**
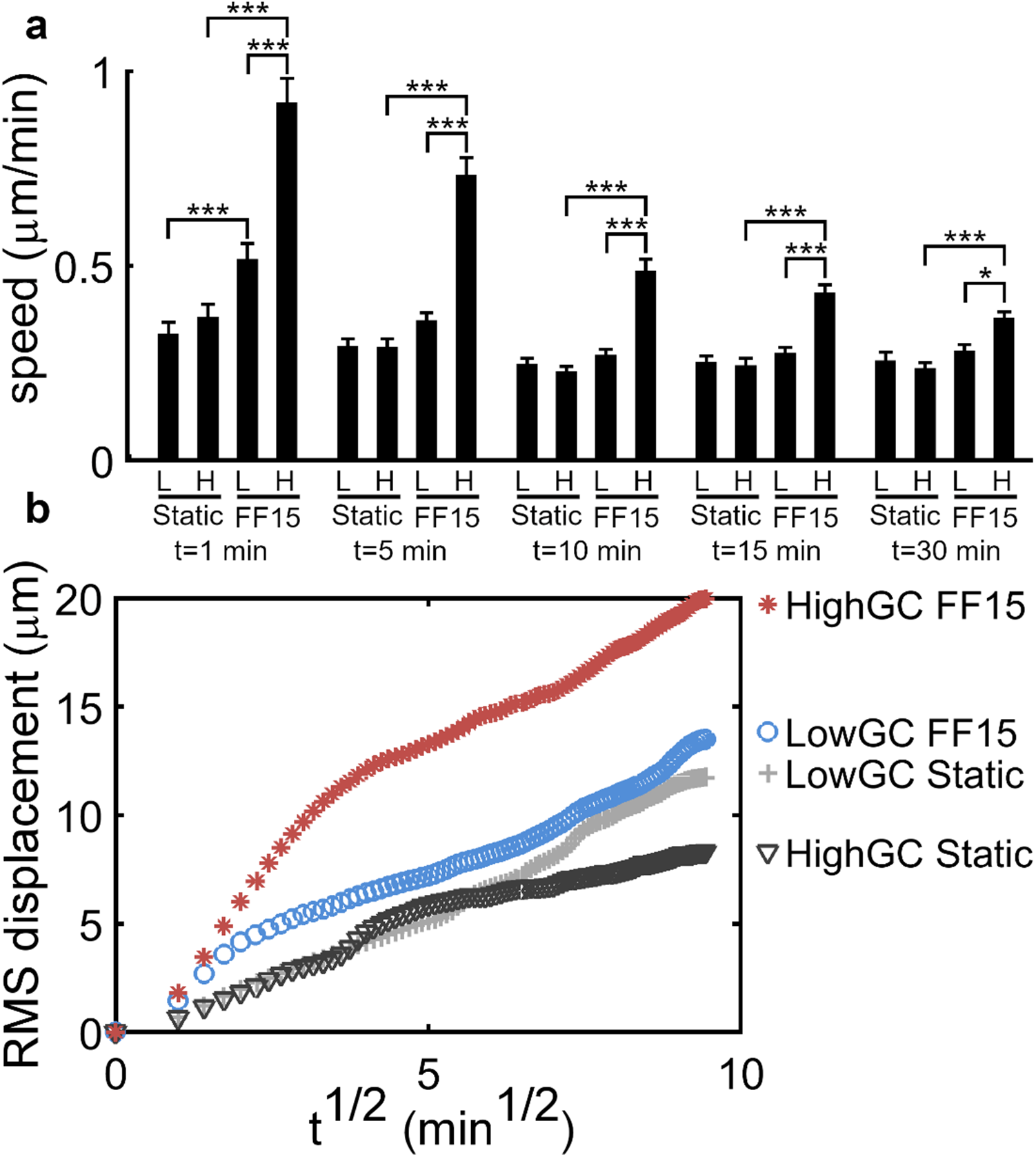
Fluid shear and high glucose promote rapid and pervasive MDA-MB-231 metastatic breast cancer cell migration. MDA-MB-231 cell migration speed (µm/min) measured for low (L) and high (H) glucose under static and flow (FF15: 15 dyne/cm^2^) conditions; t is the time in min after the fluid shear onset. Cell speed was highest right after the fluid shear onset (t = 1 min) and decreased with time. The stimulatory effects of fluid shear and high glucose were pervasive, as quantified by the overall group migration for all participating cells is shown in the RMS vs. t^1/2^ plot. Motility coefficient was LowGC Static: 1.38, HighGC Static: 0.84, LowGC FF15: 1.25, and HighGC FF15: 1.76. *, ***: p < 0.05 and 0.001, respectively. Bar graphs are plotted as mean ± SEM.

### Inhibiting FAK disrupts fluid shear-induced MDA-MB-231 TNBC cell migration under high glucose

Next, we tested the role of FAK in the migration capability of MDA-MB-231 cells under the combined effect of glucose concentration and fluid shear. The FAK inhibitor (FAK14) was applied, and MDA-MB-231 cell migration was assessed at the condition that showed the highest impact on cell migration, i.e., high glucose with fluid shear. In the raw migration tracks (Fig. 4a) and vector plots (Fig. 4b), diminished migration under FAK inhibition is clearly visible for fluid sheared MDA-MB-231 cells under high glucose. The average displacement of MDA-MB-231 cells under fluid shear and high glucose was significantly decreased by FAK14, approaching the level close to that seen with MDA-MB-231 cells under static condition (Fig. 4c, in which unflowed MCF-10A cell result in low glucose was marked as a control for comparison). Similar trends were observed for the confinement ratio (Fig. 4d) and arrest coefficient (Fig. 4e). Combined, these results show that FAK inhibition significantly abolishes the migration-promoting effects of high glucose and fluid shear on all measures of MDA-MB-231 cell migration.

**Fig. 4.**
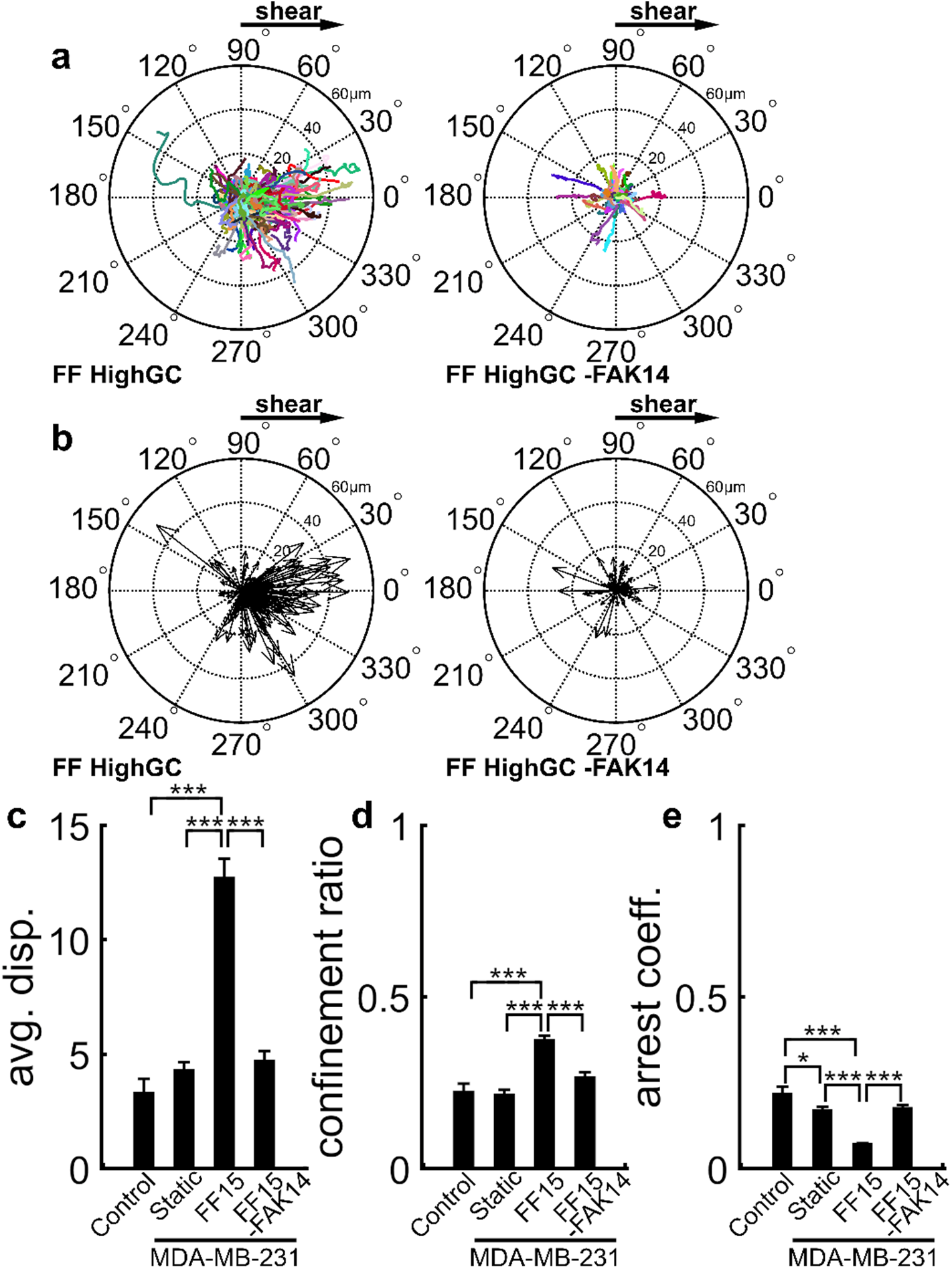
FAK inhibition abrogates the combined migration-stimulatory effect of fluid shear and high glucose in MDA-MB-231 breast cancer cells. Co-centered cell migration tracks (a) and corresponding vector plots (b) for fluid flowed cells under high glucose without or with FAK inhibitor (FAK14). Fluid shear-induced MDA-MB-231 cell migration length under high glucose was significantly suppressed by FAK14 (c) (FF15: 15 dyne/cm^2^). The migration inhibitory effects of FAK14 were also seen in confinement ratio (d) and arrest coefficient (e). Static MCF-10A cell data under low glucose were shown as controls for comparison. *, ***: p < 0.05 and 0.001, respectively. Bar graphs are plotted as mean ± SEM.

The migration speed of MDA-MB-231 cells under fluid shear and high glucose was also significantly reduced by FAK inhibition. FAK14-treated MDA-MB-231 cells in high glucose and under fluid shear had migration speeds comparable in magnitude to those seen with static cells throughout the experiment (Fig. 5a). The notable difference in migration capability is evident in the RMS displacement plot (Fig. 5b). The MDA-MB-231 cells in high glucose under fluid shear had a large ensemble-averaged group displacement with a steep slope, whereas the static and FAK-inhibited flow groups under high glucose had similar trends of significantly impaired ensemble average migration, thus demonstrating a significant impairment due to FAK inhibition. The motility coefficient was 1.76 for MDA-MB-231 cells in high glucose under fluid shear. On the other hand, cells in high glucose under static condition had a motility coefficient of 0.84, comparable with FAK inhibitor-treated high glucose and fluid shear group with a value of 0.86. This again reveals that fluid shear-stimulated MDA-MB-231 cell migration under high glucose is suppressed by FAK inhibition down to the level of MDA-MB-231 under static condition. Further, although we found that fluid shear stress and hyperglycemic condition both promoted migration, neither stimuli could reverse the effects of FAK inhibition. These results support that FAK is critical to govern cell migration capability of TNBC cells promoted by mechanical (fluid shear) and metabolic (glucose concentration) factors.

**Fig. 5.**
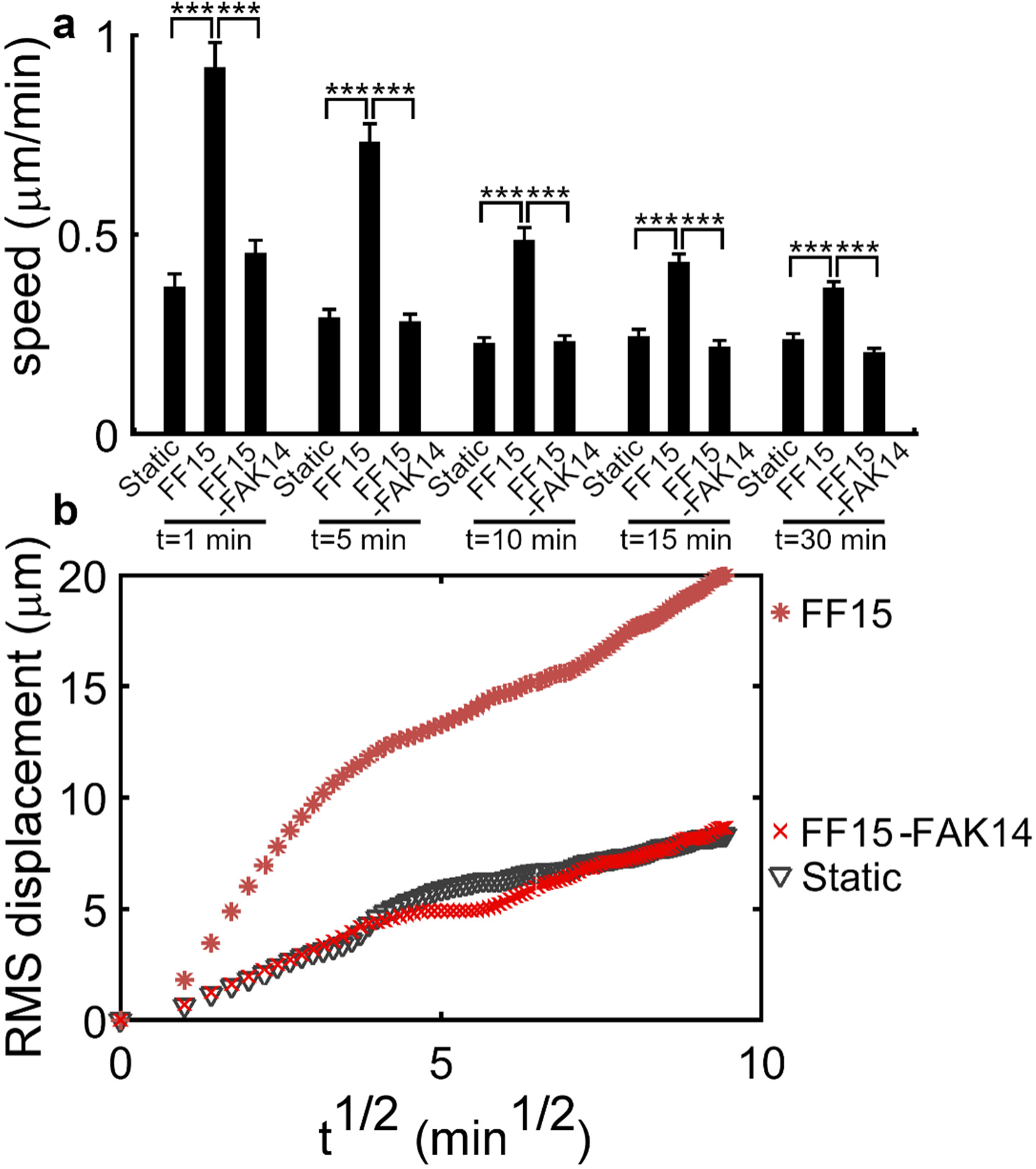
Stimulations in migration speed and ensemble averaged migration of MDA-MB-231 cells by high glucose and fluid shear are inhibited by FAK inhibition. FAK inhibition by FAK14 significantly decreased MDA-MB-231 cell migration speed under fluid shear (FF15: 15 dyne/cm^2^) and high glucose (a); t is the time in min after the fluid shear onset. Overall migration of the group was also blocked by FAK14 as in the RMS vs. t^1/2^ plot (b). Motility coefficient was Static: 0.84, FF15: 1.76, and FF15-FAK14: 0.86, all under high glucose. ***: p < 0.001, Bar graphs are plotted as mean ± SEM.

### FAK control of MDA-MB-231 migration in scratch wound and Boyden chamber assays

To complement our findings above, we also measured the migration capability of MDA-MB-231 cells using the scratch wound healing assay. We found comparable results in the closure of the scratch wound as well as the number of cells migrating through the scratch wound under low and high glucose conditions (Fig. 6a,b,c). On the other hand, inhibiting FAK by FAK14 significantly suppressed the wound closure measured at 24 h under both low and high glucose conditions (Fig. 6a,b). The number of cells migrating through the scratch area was also significantly decreased by FAK inhibition under low and high glucose concentrations (Fig. 6c).

**Fig. 6.**
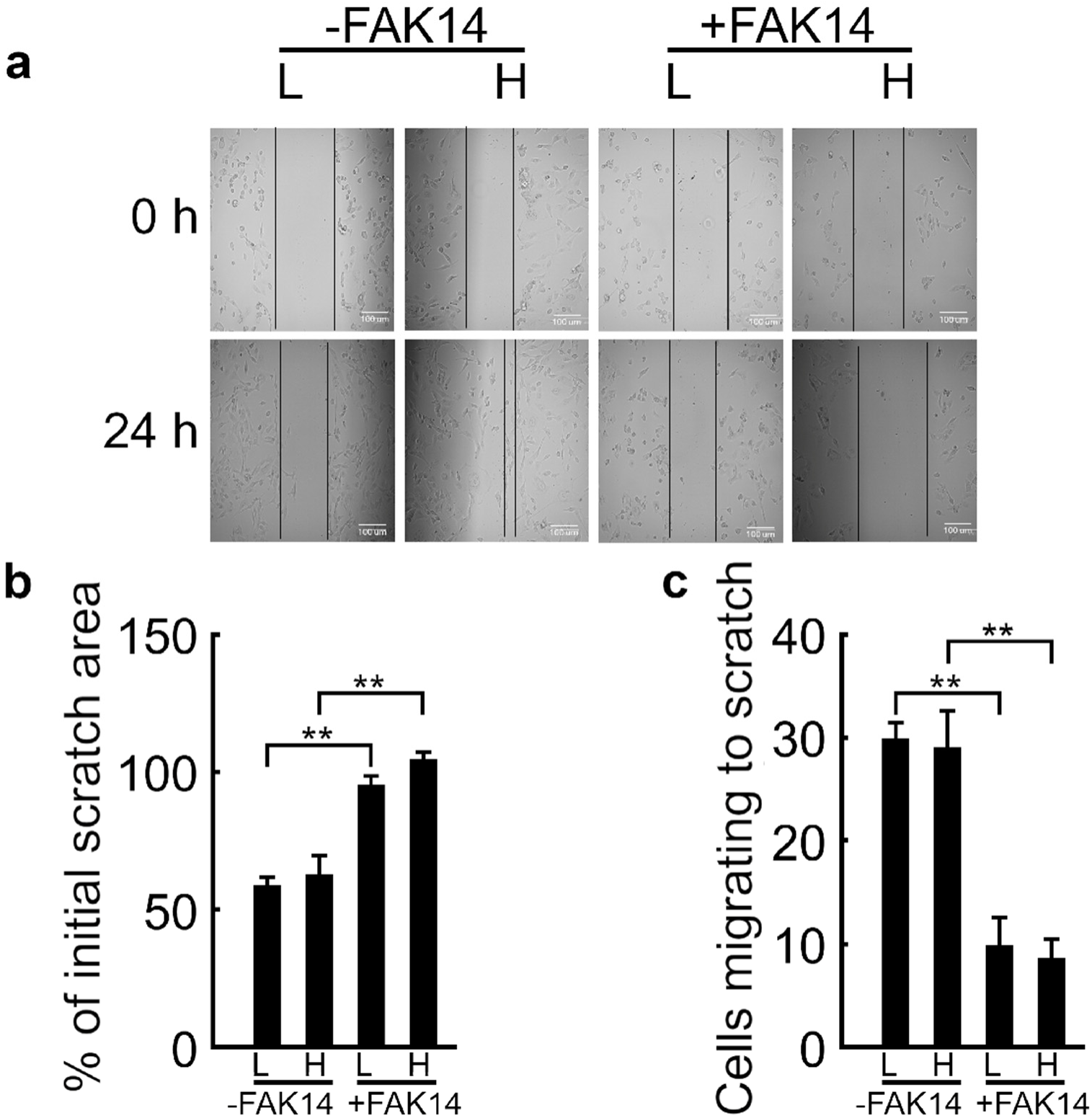
FAK inhibition prevents the migration of MDA-MB-231 cells towards the scratch wound area. (a) Initial scratch area at 0 h and the area imaged at 24 h under low (L) and high (H) glucose without or with FAK inhibitor (FAK14). There was no appreciable difference in scratch wound area with low (L) and high (H) glucose conditions. On the other hand, FAK14 considerably suppressed the covering of the scratch (b) and movement of cells towards the scratch (c). **: p < 0.01 (n = 5). Bar graphs are plotted as mean ± SEM.

The migration of MDA-MB-231 cells was also evaluated using Boyden chambers, where the ability of cancer cells to move across an artificial membrane towards serum factors was monitored. The number of cells migrated after 24 h was not significantly different between low and high glucose conditions, but was reduced by about 50% in the presence of FAK inhibitor (Fig. 7a,b). Overall, the combined results support the conclusion that while TNBC cell migration is critically dependent on FAK under all conditions, high glucose selectively enhances the FAK-dependent TNBC cell migration under fluid shear stress environment.

**Fig. 7.**
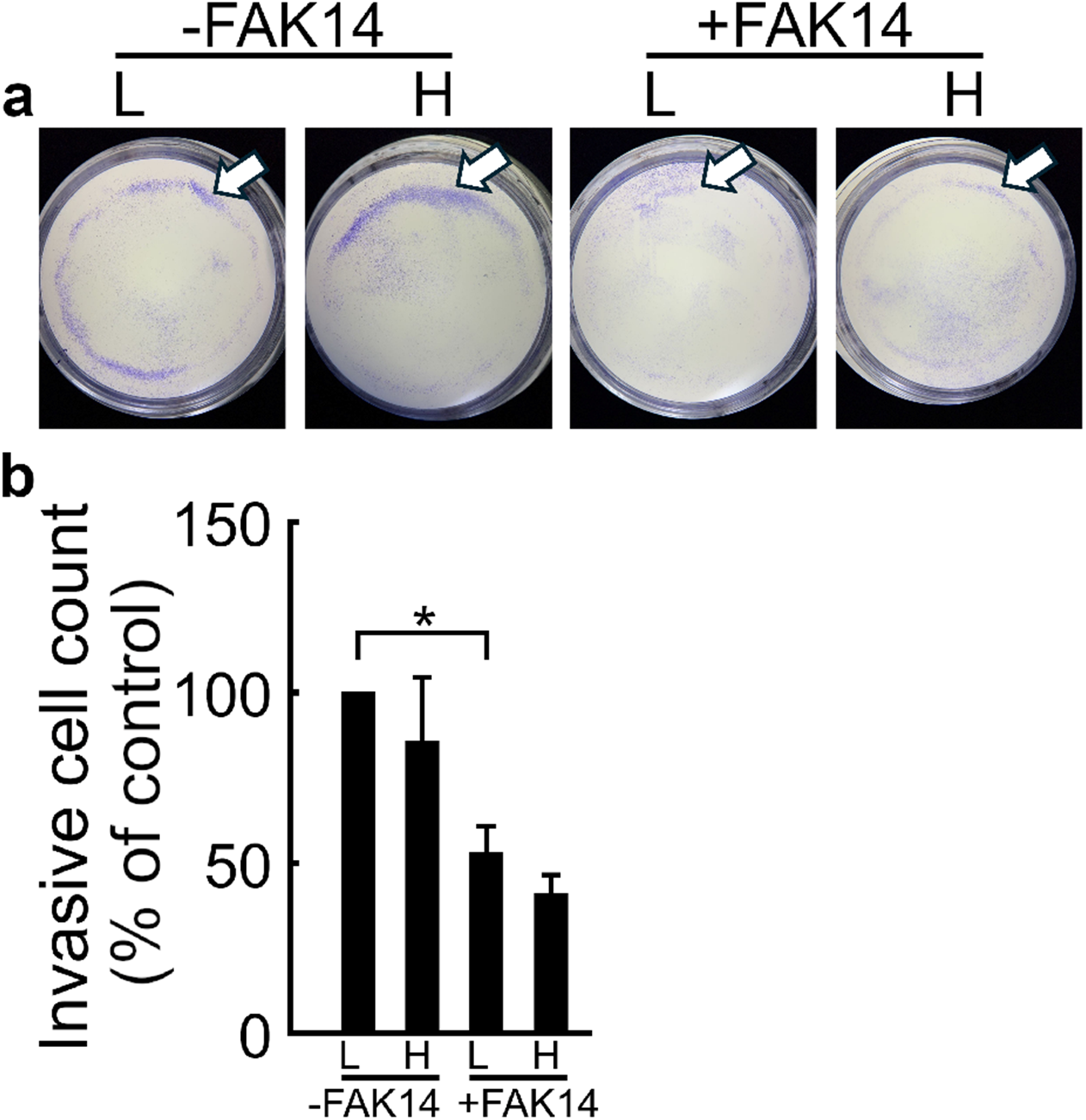
FAK inhibition prevents the transwell migration of MDA-MB-231 cells. (a) Images of transwell migration assay for MDA-MB-231 cells under low (L) and high (H) glucose without or with FAK inhibitor (FAK14). Arrows example the regions of stained cells. (b) The invasive cell count was not affected by glucose concentration, while FAK14 substantially decreased cell movement across the membrane. Invasive cell count is presented as % of low glucose control without FAK14. *: p < 0.05 (n = 5). Bar graphs are plotted as mean ± SEM.

## Discussion

The higher mortality of breast cancer patients with concurrent T2DM necessitates mechanistic dissection of links between high glucose, a feature of T2DM patients, and the metastatic process. Cell migration is a critical component of the metastatic program. TNBC is a particularly aggressive subtype of breast cancer that is highly prone to metastasize and currently lacking in targeted therapies. Here, we used the highly metastatic TNBC cell line, MDA-MB-231, to understand the impact of high glucose on the motility of metastatic breast cancer cells under static vs. fluid shear conditions to better elucidate the impact of diabetes-associated hyperglycemic state on the TNBC behavior. Our findings reveal that high glucose selectively accentuates the motility of metastatic TNBC cells under fluid shear stress, supporting the idea that metabolic and mechanophysical cues may combine to promote the pro-metastatic behavior in TNBC cells. Our findings further show that FAK is required for the high glucose-promoted migration of MDA-MB-231 cells under fluid shear, supporting the potential of targeting FAK to reduce the collaborative promotion of TNBC metastasis through hyperglycemia and mechanophysical factors impinging on tumor cells.

While it is becoming increasingly recognized that breast cancer patients with coexistent diabetes exhibit a significantly higher risk of mortality compared with those without diabetes [2], the exact underlying mechanisms are yet to be fully revealed. We provide direct evidence that highly metastatic MDA-MB-231 TNBC cells display significantly stimulated migratory capability under the diabetes-relevant high glucose condition when specifically exposed to the physiologically relevant fluid flow-induced shear stress milieu (Figs. 2,3). Such a fluid shear dependency of metastatic MDA-MB-231 cell migration was in contrast to that of normal epithelial MCF-10A cells displaying increased migration by fluid shear under both low and high glucose conditions (Fig. 1). In the presence of FAK inhibitor, fluid shear-triggered MDA-MB-231 cell migration under high glucose condition was significantly suppressed (Figs. 4,5), suggesting the regulatory role of FAK in metastatic breast cancer cell sensing of the mechanical (fluid shear stress) as well as biochemical (glucose concentration) cues during migration. In the scratch wound healing and Boyden chamber transwell migration assays (in which fluid shear was not applied), differences in MDA-MB-231 migration between low and high glucose conditions did not reach statistical significance; on the other hand, FAK inhibition clearly impaired the MDA-MB-231 cell migration under both low and high glucose concentrations (Figs. 6,7).

Breast cancer cells encounter fluid shear stresses starting from the primary tumor sites all the way to metastatic target tissues. An increased pressure at the tumor site due to tumor expansion causes an interstitial flow ranging from about 0.1 to 12 dyne/cm^2^ shear stresses [24]. As they travel through circulation, cancer cells are exposed to blood flows producing shear stresses of about 6 to 70 dyne/cm^2^, depending on venous or arterial components of the circulatory system [25]. The metastasis sites, for instance, bone (one of the primary metastasis sites for breast cancer cells), provides interstitial fluid shear stresses of about 0.06 to 30 dyne/cm^2^ within the bone tissue due to the three-point bending motion of the bone [26]. Our current data obtained at 15 dyne/cm^2^ shear stress, within the range of shear encountered in vivo, provide clear evidence that metastatic MDA-MB-231 breast cancer cells display distinct fluid shear sensitivity by exhibiting significantly increased migration distance, confinement ratio, number of cells moving with the flow, cell speed, ensemble average RMS displacement, and motility coefficient as well as decreased arrest coefficient (Figs. 2,3). Intriguingly, such a fluid shear sensitivity of MDA-MB-231 TNBC cells was observed under the high glucose condition, but at a marginal degree under low glucose. This may have a direct implication on the diabetes-cancer crosstalk by pointing to the potential of the hyperglycemic state to promote the metastatic TNBC cell migration in response to the physiological fluid shear environment. We presume that enhanced migration of TNBC cells under high glucose condition and fluid shear stress environment could play a role in increasing metastasis in TNBC patients with concurrent T2DM, requesting future in vivo studies to test this idea.

Our FAK inhibitor data suggest a mediatory role of FAK in metastatic breast cancer cell migration under biochemical-mechanical combinatory conditions. Specifically, the result that fluid shear-induced MDA-MB-231 migration under high glucose was significantly impaired by the FAK inhibitor (Figs. 4,5) suggests that FAK-mediated focal adhesion could play a vital role in such breast cancer cell migration. For achieving migration, cells need to form new focal adhesions at the leading edge of the migrating cell while disassembling those at the trailing edge [7]. Cells can recruit FAK at the new adhesion sites during migration to reinforce focal adhesion and to further link to the cytoskeleton [10]. Our previous work with mesenchymal stem cells (MSCs) with stable silencing of FAK via small hairpin RNA (shRNA) demonstrated that FAK is required for MSC migration under fluid shear [27]. The current data indicate that a similar cascade acts for breast cancer cell migration, with FAK inhibition likely suppressing the capability of TNBC cells to create new attachment sites required for migration. Importantly, the data in this study provide new insights that high glucose condition could promote the fluid shear sensitivity of MDA-MB-231 cell migration and that the stimulatory impact of high glucose-fluid shear is impaired by FAK inhibition. This implies a vital role of FAK in metastatic breast cancer cell migration under the combined influence of biochemical (glucose) and mechanical (flow) cues promoting cell migration. FAK is recently tested as a potent therapeutic target for breast cancer treatments [11–14]. FAK has been also proposed to be involved in the evolution of diabetes and related diseases [21,22]. However, much remains unknown on the regulatory role of FAK in diabetic metastatic triple negative breast cancer. Our FAK inhibitor results support the idea that FAK is required for high glucose-fluid shear-induced TNBC cell migration and propose a multi-factor-responsive role of FAK to regulate breast cancer progression for diabetic patients. In-depth future studies of this molecular crosstalk are expected to reveal important insights into the role of FAK and associated avenues to target in breast cancer patients with T2DM.

In traditional scratch wound and Boyden chamber assays, high and low glucose conditions did not result in significant changes in MDA-MB-231 migration. This might be potentially due to the fact that physiologically relevant fluid shear environments are not embedded in those assays.

On the other hand, the FAK inhibitor was quite effective in suppressing the mobility of MDA-MB-231 cells in these conventional assays (Figs. 6,7), again highlighting the role of FAK in breast cancer cell migration. We have previously observed that diabetic states increase the MDA-MB-231 cell proliferation, which was mediated through the polyamine metabolic pathway [28]. In relation to this study, we recently evidenced the role of cytoskeletal regulatory kinases, such as FAK and ROCK, in the cytoarchitecture of diabetic breast cancer cells [29]. We showed that diabetes-relevant high glucose media condition altered the cytoskeletal architecture of breast cancer cells, thereby decreasing the elasticity of the cells. Further, inhibitors of FAK and ROCK diminished such effects of high glucose. In combination with current data demonstrating FAK-mediated breast cancer cell migration under high glucose and fluid shear (Figs. 4,5), these indicate the crucial role that FAK plays in regulating the cytoskeletal architecture and its potential outcome, such as tumor cell migration, under diabetic and fluid shear situations. Altogether, the impacts of the hyperglycemic condition on increased proliferation along with stimulated migration via FAK mechanism suggest a FAK-mediated collaborative signaling by biochemical and mechanical cues as an underlying mechanism of how diabetes may increase breast cancer progression and metastasis.

Future studies on varying flow shear milieus will provide more knowledge. Different fluid shear stress regimens, e.g., shear stress magnitudes recapitulating those at primary tumor site, blood vessel, and metastasis site, could reveal the extent to which breast cancer cells alter their migration behavior as a function of shear stress level. Varying fluid flow modes, such as steady, pulsatile, or cyclic and 2D monolayer flow vs. 3D perfusion flow, may also affect breast cancer cell migration, as we have discussed previously [30]. While our studies on breast cancer cells (this and previous study [23]), MSCs [27], and osteoblasts [31] show that cell migration under fluid shear follows along the flow direction, a study by Polacheck et al. [32] evidenced that breast cancer cells may migrate with or against the flow direction depending on the cell seeding density. They showed that breast cancer cells migrate with the flow direction when seeded at a low density but against the flow at a high density. Taken together, more systematic, extensive studies with altering cell culture conditions to represent varying in vivo environments will provide more in-depth knowledge on the regulation of breast cancer cell migration during metastatic processes.

In summary, this study assessed the effect of the diabetes-related hyperglycemic condition and physiological fluid shear stress environment on the migration of highly metastatic TNBC cells, MDA-MB-231. Our data demonstrate that high glucose and fluid shear collaboratively accentuate TNBC cell migration in a FAK-dependent manner, supporting a mechanism on the impact of T2DM on breast cancer outcomes and the exploitation of FAK as a potential therapeutic target to deal with biochemical-mechanophysical combined impacts.

## Materials and Methods

### Reagents and supplies

Cell lines (MDA-MB-231, MCF-10A), Eagle’s minimum essential medium (EMEM), and fetal bovine serum (FBS) were obtained from American Type Culture Collection (ATCC, Manassas, VA, USA). Dulbecco’s Modified Eagle Medium/F12 (DMEM/F12, 1:1), phenol red-free DMEM, horse serum, penicillin/streptomycin, GlutaMAX, and PrestoBlue cell viability reagent were obtained from Thermo Fisher Scientific (Rockford, IL, USA). Epidermal growth factor (EGF) was obtained from Peprotech (Rocky Hill, NJ, USA). Glucose, insulin, hydrocortisone, and cholera toxin were obtained from Millipore Sigma (Milwaukee, WI, USA).

### Cell culture conditions

MDA-MB-231 TNBC cells and MCF-10A immortal normal human mammary epithelial cells were cultured at 37°C in a humidified atmosphere with 5% CO2, under the following conditions. Each cell line was maintained in media conditions standard for the specific cell line before being transferred to the flow media. MDA-MB-231 cells were grown in DMEM/F12 supplemented with 5% FBS, 1% GlutaMax, 100 IU/ml penicillin, and 100 μg/ml streptomycin. MCF-10A cells were grown in DMEM/F12 with 5% horse serum, 20 ng/ml EGF, 0.5 µg/ml hydrocortisone, 100 ng/ml cholera toxin, 10 μg/ml insulin, 100 IU/ml penicillin, and 100 μg/ml streptomycin. After reaching assay-specific confluency, cells were cultured for 48 h in glucose-free, phenol red-free DMEM media supplemented with varying concentrations of glucose (5 mM or 25 mM), 1% FBS/1% horse serum, 1% GlutaMAX, 100 IU/ml penicillin, and 100 µg/ml streptomycin.

### Cell culture for fluid shear tests

Once cells reached 80% confluence in a petri dish, cells were collected and seeded at 1:12 ratio on 25×75 mm^2^ glass slides using 1 mL media. Cells were allowed to attach for 90 min on the glass slide before additional media was added to the plate. After 24 h, the medium was changed to glucose starvation media for 1 h, and then replaced with either low (5 mM) or high (25 mM) glucose media. Fluid shear experiments were conducted after 48 h exposure to high and low glucose media. MCF-10A cells were cultured and treated using the same protocol and timeline as described above. Fluid flow media with reduced serum for these cells consisted of DMEM supplemented with 2.5% serum and 1% penicillin-streptomycin. For FAK inhibition studies, slides were placed in media containing 10 μM FAK inhibitor 14 (FAK14, Santa Cruz, sc-203950, Dallas, TX, USA) for 1 h. Then, slides were rinsed in phosphate buffered saline (PBS) before being placed in the flow chamber. The exposure to the selected FAK14 concentration did not induce cytotoxicity as assessed by the PrestoBlue cell viability reagent.

### Fluid shear stimulation of cells and time lapse imaging

Fluid flow was generated using the Flexflow flow device (Flexcell International, Burlington, NC, USA). A steady fluid flow at 15 dyne/cm^2^ shear stress (labeled FF15) was applied to the cells for 90 min with 1 min time lapse imaging interval, following our published protocols [23,27,31]. The fluid sheared cell samples were compared with the static control in which the cell-seeded slides were placed in the same flow chamber but not exposed to flow. The flow system was set up using the standard components from the manufacturer in which Masterflex L/S16 tubing connects the media reservoir, two pulse dampeners, and the Flexflow flow device. The flow system was sterilized with 70% ethanol and flushed twice with de-ionized water before flow. After cleaning and flushing, the flow media was added to the reservoir and bubbles were purged from the flow loop. A leak test was performed before placing the device on the microscope for time lapse imaging. Flow media was maintained at 37°C in a water bath. A Leica DMI 4000 microscope was used to capture phase contrast images of the cells in the fluid shear chamber once every min for 90 min.

### Quantification of cell migration tracks

Our established protocols [23,27,31] were used to segment and process time lapse cell images. Minor adjustments to correct for device drift were conducted using the template matching plugin for FIJI-ImageJ [33]. After stabilization, binary masks were created for each cell using Sobel edge detection with post processing to fill in holes. Manual inspection and correction in ImageJ were used to ensure that masks fit the cell outlines and that adjacent binary masks did not overlap. The binary cell masks were tracked in time lapse analyzer (TLA) [34]. The obtained cell migration tracks were processed in our custom MATLAB script [23,27,31] to quantitatively assess cell migration via average cell displacement length, confinement ratio (how straight cells migrate), arrest coefficient (how often cells stop during migration), percentage of cells moving along the flow direction, cell migration speed, RMS displacement (ensemble-averaged cell migration from all participating cells), and motility coefficient from the RMS plot. The RMS displacement is calculated as *X* (*t*) *RMS*:

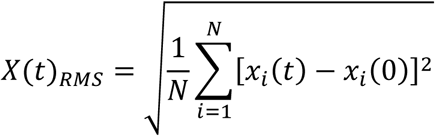

where *N* is the total number of observed cells, *xi* (*t*) is the position of the *i*-th cell at time *t*, and *xi* (0) is the position of that cell at time 0. As the RMS displacement is a good measure of group (ensemble) migration, we also refer to RMS displacement as the ensemble average throughout the paper. The motility coefficient was obtained as the slope of the RMS displacement curve with respect to the square root of time. The motility coefficient delivers the same measure as with the diffusion coefficient of the first-order diffusion kinetics [35], presenting the strength of migration of all participating cells. From the obtained time lapse images, individual cells that remained in the imaging window and did not contact other cells were analyzed for cell migration.

### Scratch wound healing assay

Cells were seeded in a 6-well plate and allowed to adhere for 24 h. After cells reached about 90% confluence, a scratch was made along the entirety of each well using a sterile tip. The medium was aspirated immediately after scratching, and the wells rinsed with PBS. The cells were subsequently treated with the indicated concentrations of glucose (5 mM or 25 mM) and FAK inhibitor. Images were taken at 0 h and 24 h after scratch was performed. Percent wound closure was then measured.

### Transwell migration assay

Cells were counted and plated at 350,000 cells/well on the top of Boyden chambers in a 6-well plate in the treatment media (no serum), with complete media containing serum added in the bottom of each chamber to assess cell migration. Cells were incubated for 24 h under indicated glucose concentrations (5 mM or 25 mM) and FAK inhibitor. The chambers were washed twice with PBS, and the migrated cells were fixed in methanol at room temperature for 15 min. Cells were stained with 0.5% crystal violet for 30 min, washed with water to remove excess color. The membranes were viewed under a stereoscope and images taken. Cell counts were analyzed using ImageJ.

### Statistical analysis

All values obtained were expressed as mean ± standard error of mean (SEM). The results shown were analyzed with Graph Pad software (Prism 5.0) and are representative of at least three independent experiments. Statistical comparison between more than two distinct experimental groups was performed using one-way ANOVA followed by Tukey’s post hoc test. Cell migration data that violated the statistically normal distribution assumption of ANOVA were transformed into log10, tested, and the results were back transformed.

## Data availability

The dataset generated during the study is available from corresponding authors upon reasonable request.

## Acknowledgments

The authors thank the funding supports from University of Nebraska Collaboration Initiative Seed Grants (D.D., S.C., and J.Y.L.; S.C. and J.Y.L.; H.B. and J.Y.L.); Nebraska Center for Integrated Biomolecular Communication (NIH/NIGMS P20GM113126, J.Y.L.); Nebraska Center for the Prevention of Obesity Diseases (NIH/NIGMS P20GM104320, J.Y.L.); University of Nebraska-Lincoln Biomedical Research Seed Grants (J.Y.L.); NE DHHS Stem Cell Research Project (2023-04, J.Y.L); National Institute for General Medical Science (NIH/NIGMS 5R35GM150504, S.C.). Cell culture and fluorescence microscopy facilities used in this project were supported by grants from the National Center for Research Resources (NIH/NCRR 5P20RR016469) and the National Institute for General Medical Science (NIH/NIGMS 8P20GM103427 and 5P20GM103427).

## Author contributions

B.D.R., S.C. (Choi), H.B., L.M., D.D., S.C. (Chandra), and J.Y.L. conceived and designed the study. B.D.R., E.K., T.B., O.O., and S.V-D. performed the experiments. B.D.R. and S.C. (Chandra) provided figures from experimental data. B.D.R., S.C. (Chandra), and J.Y.L interpreted the results and performed discussions. B.D.R. and S.C. (Chandra) wrote the initial manuscript. B.D.R., S.C. (Choi), H.B., L.M., D.D., S.C. (Chandra), and J.Y.L. revised the manuscript. S.C. (Chandra) and J.Y.L. supervised the project.

